# Molecular basis for *B. pertussis* interference with complement, coagulation, fibrinolytic and contact activation systems: The cryo-EM structure of the Vag8-C1 inhibitor complex

**DOI:** 10.1101/2020.10.05.327577

**Authors:** Arun Dhillon, Justin C. Deme, Emily Furlong, Dorina Roem, Ilse Jongerius, Steven Johnson, Susan M. Lea

## Abstract

Complement, contact activation, coagulation, and fibrinolysis are serum protein cascades that need strict regulation to maintain human health. Serum glycoprotein, C1-inhibitor (C1-INH) is a key regulator (inhibitor) of serine proteases of all the above-mentioned pathways. Recently, an autotransporter protein, Virulence Associated Gene 8 (Vag8) produced by the whopping cough causing pathogen, *Bordetella pertussis* has been shown to bind and interfere with C1-INH function. Here we present the structure of Vag8: C1-INH complex determined using cryo-electron microscopy at 3.6 Å resolution. The structure shows a unique mechanism of C1-INH inhibition not employed by other pathogens where Vag8 sequesters the Reactive Centre Loop of the C1-INH preventing its interaction with the target proteases.

**Importance:** The structure 105 kDa protein complex is one of the smallest to be determined using cryo-electron microscopy at high resolution. The mechanism of disrupting C1-INH revealed by the structure is crucial to understand how pathogens by producing a single virulence factor can disturb several homeostasis pathways. Virulence mechanisms such as the one described here assume more importance given the emerging evidence about dysregulation of contact activation, coagulation and fibrinolysis leading to COVID-19 pneumonia.

## Introduction

Protein cascades coordinate key processes for health within human serum, in particular immune and inflammatory responses (complement and contact activation) and control of clotting (contact activation, coagulation and fibrinolysis) (1–3). Although independent processes, coordination between the pathways occurs by shared regulation, particularly by C1-inhibtor (C1-INH) (4). C1-INH inhibits serine proteases involved in activation and control of these systems by formation of protease-C1-INH complexes such that the level of these complexes is directly proportional to the level of *in vivo* activation of all four systems (5). C1-INH is established as a key regulator of complement via inhibition of the activation proteases C1r, C1s, MASP-1 and MASP-2 and is the dominant inhibitor of plasma kallikrein (contact activation system), coagulation factors XIIa and XIa and thrombin (6–10). The mode of inhibition of these proteases involves interaction between the Reactive Centre Loop (RCL) of the C-terminal Serpin domain of C1-INH to form a covalently linked acyl-enzyme complex that distorts the enzyme active site and is irreversibly bound (11–13). Additionally, C1-INH has been implicated in regulation of fibrinolysis via action against tissue-type plasminogen activator (tPA) and plasmin – although study of this is complicated by the fact that both these enzymes will also cleave C1-INH (14, 15).

Whooping cough (pertussis) is an infectious disease of the respiratory system caused by the Gram-negative bacterium *Bordetella pertussis* (16). *B. pertussis* employs a range of virulence factors to colonise the human host and evade immune responses (17). Some of these factors e.g. Virulence associate gene 8 (Vag8), *Bordetella* Resistance to Killing A (BrkA), Filamentous hemagglutinin (FHA) and B. pertussis autotransporter protein C (BapC) have been implicated in evasion of the complement system (18–21). While the mechanisms of action of BrkA, BapC, FHA are still unclear, Vag8, a 95 kDa auto-transporter protein was recently shown to interfere with the complement and contact systems by binding to C1-INH leading to bacterial complement evasion (22, 23). Auto-transporters represent the type V bacterial secretion system and possess a C-terminal membrane embedded β-barrel domain that facilitates the translocation of the N-terminal passenger domain, responsible for effector functions, across the outer membrane (24). In case of Vag8 the cleaved N-terminal domain has been detected in bacterial culture supernatant in addition to the full length Vag8 being presented on outer membrane vesicles (OMVs), and on the cell surface (22). Deletion of the gene encoding Vag8 predisposes *B. pertussis* to complement mediated killing (18, 22).

Although C1-INH is an inhibitor of complement activation, targeting C1-INH activity is used as a strategy for complement evasion by a range of different pathogens. *Streptococcus pyogenes*, and *Legionella pneumophila* use enzymes, SpeB and ChiA respectively, to cleave C1-INH (25, 26), while *Plasmodium falciparum, Borrelia recurrentis and Salmonella typhimurium* depend on *Pf*MSP3.1, CihC, and lipopolysaccharide (27–29), respectively to capture C1-INH on the cell surface. A hybrid of the above two strategies of C1-INH targeting has been proposed to be used by *E. coli O157:H7* involving capture of C1-INH on the cell surface followed by an enzymatic cleavage (30). Whilst targeting an inhibitor to the pathogen surface is a self-evident way of enhancing immune evasion, the utility of destruction of C1-INH is less obvious but occurs due to the fact that removal of C1-INH from serum leads to rapid, catastrophic activation of complement, leading to depletion of activity and so, perversely, less complement attack on the pathogen (22).

Globally, pertussis is responsible for a large number of infant deaths, especially in low income countries and is a financial burden even in developed economies (31, 32). Despite extensive vaccination programs *B. pertussis* infections are on the rise again (33). Reasons to explain the rising infections have been contentious and include waning of immunity generated by acellular pertussis vaccines, and evolution of more pathogenic strains (34–37) therefore, a molecular understanding of the mode of action of *B. pertussis* virulence factors such as Vag8 is desirable. More broadly, with evidence mounting that activation of coagulation and excessive cytokine release are key drivers of COVID-19 pneumonia and mortality with contact activation appearing particularly important in driving pathologic upregulation of inflammatory mediators and coagulation, interest in pathogenic mechanisms acting on these systems is further increased (38–41).

To that end, we have determined the structure of the Vag8 passenger domain in complex with the C1-INH Serpin domain using single particle cryo-electron (cryo-EM) microscopy to 3.6 Å resolution. The Cryo-EM structure of this complex reveals that Vag8 non-covalently sequesters the reactive centre loop (RCL) of C1-INH in the groove of the elongated passenger domain so preventing C1-INH/protease interactions and regulation. Thus *B. pertussis* overrides complement regulatory control by a unique mechanism not previously seen in other pathogens. Sequestration of C1-INH in this manner not only leads to complement evasion but also promotes kallikrein activation, leading to increased levels of the vasoactive bradykinin, increased fibrinolysis, and coagulation. Thus *B. pertussis* widely perturbs serum activities across a broad spectrum by production of a single protein molecule.

## Results

To better understand how *B. pertussis* subverts C1-INH function we heterologously expressed and purified both the passenger domain of Vag8 and the Serpin domain of C1-INH (Figure 1a, b). When mixed at an approximately equimolar ratio the proteins formed a complex that could be separated from a small amount of residual isolated C1-INH by size-exclusion chromatography (Figure 1a, b). This Vag8:C1-INH complex was then concentrated to 0.5 mg/ml and applied to Quantifoil R1.2/1.3 carbon-coated grids before blotting using a Mark IV Vitrobot and plunge freezing in liquid ethane. Single particle cryoEM data were collected using a Titan Krios at 300kV equipped with a Gatan BioQuantum and K3 detector as described in the methods. The small size of the complex (∼100 kDa) meant that individual particles were difficult to discern at the micrograph level (Figure 1c), however manual picking of ∼1000 particles followed by 2D classification generated 2D averages that were used for automated picking of more than 40,000 movies, collected from three grids (Figure 1d). Data were processed as shown in the workflow (Figure 1d) using both SIMPLE3.0 (42) and RELION3.1 (43) to yield a final volume based on 687,883 particles with an estimated resolution (by gold-standard FSC, 0.143 criterion) of 3.6 Å (Figure 1e). Calculation of a local resolution filtered volume (Figure 1f; RELION 3.1, (43)) demonstrates that the core of Vag8 and size of interaction with C1-INH is well defined, with a resolution estimate of 3.5Å despite the small size of this complex placing it amongst the ten smallest structures determined to date using this method (44).

**Figure 1.**
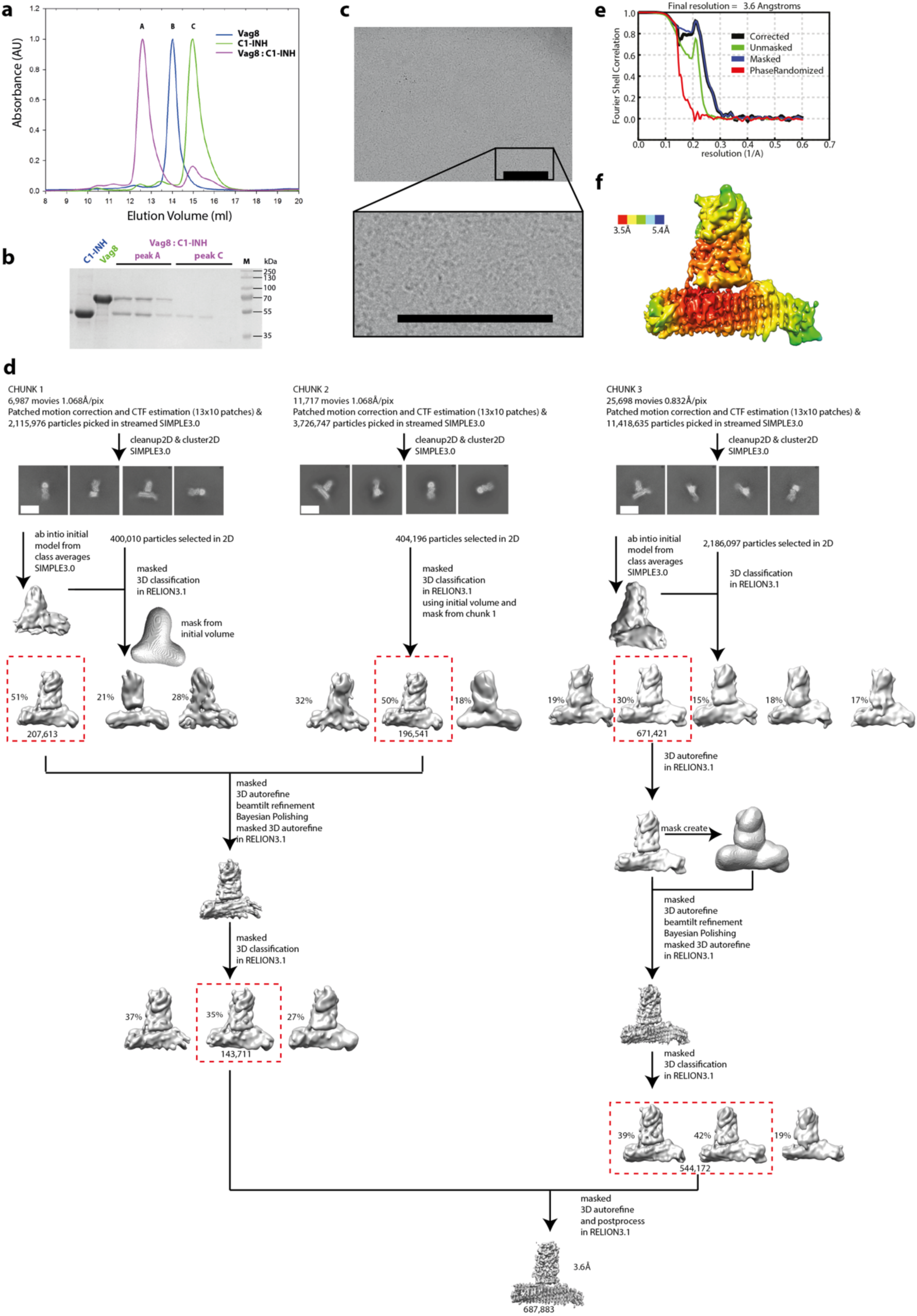
Determination of the single particle CryoEM structure of the Vag8:C1-INH complex at 3.6 Å. (**a**) Size exclusion chromatography analysis shows that Vag8 binds C1-INH to form a complex (purple). 100µl of an approximately 1:2 molar ratio of Vag8 : C1-INH were mixed and purified using a S200 increase chromatography column. A, B and C denote the locations at which Vag8:C1-INH complex, Vag8 and C1-INH respectively elute. (**b**) Fractions under peaks A & C from Vag8:C1-INH purification when run on 15% (w/v) SDS-PAGE gel confirm that the peak at location A contains Vag8 bound to C1-INH while unbound C1-INH elutes in peak C. (**c**) A representative micrograph of Vag8:C1-INH complex on a carbon-coated grid. Scale bar 200 Å (**d**) cryo-EM data of Vag8:C1-INH complex were collected and initially processed as 3 different chunks (Chunk 1, 2 and 3) and combined at later stages during processing using SIMPLE 3.0 and RELION 3.1. Masked 3D classification of chunk 2 data was done using the initial volume and mask from chunk 1. Subsequently, selected particles from chunk 1 and 2 were combined and masked 3D classification was performed. Selected particles from this data set were combined with selected particles from chunk 3 data obtained after masked 3D auto-refine and masked 3D classification. This final combined data set was then auto-refined and postprocessed in RELION 3.1 resulting in 3.6 Å volume. Scale bar on 2D averages is 50 Å (**e**) Gold-standard Fourier Shell Corelation curves of Vga8:C1-INH complex volumes postprocessed in RELION 3.1. Curves: red, phase-randomized; green, unmasked; blue, masked; black, corrected (**f**) Volume coloured by estimated Local resolution (Å).

A *de novo* model was built manually using program COOT (45) for the region 54-481 of Vag8. Although residual density could be seen in the volume both N- and C-terminal to this region (Figure 2a), it was not possible to build an atomic model for residues 40-53 and 482-610. The model of the active form of the C1-INH Serpin domain (46) was placed and remodelled to fit the volume, with the only major changes in conformation being within the RCL which is seen to be sequestered within the cleft of the Vag8 beta-barrel fold. Figure 2b shows the quality of the volume around key-side chains within the binding site. Following further cycles of manual rebuilding and real-space refinement in PHENIX (47) lead to the generation of the model presented in Figure 2 and described in Table 1.

**Table 1.**
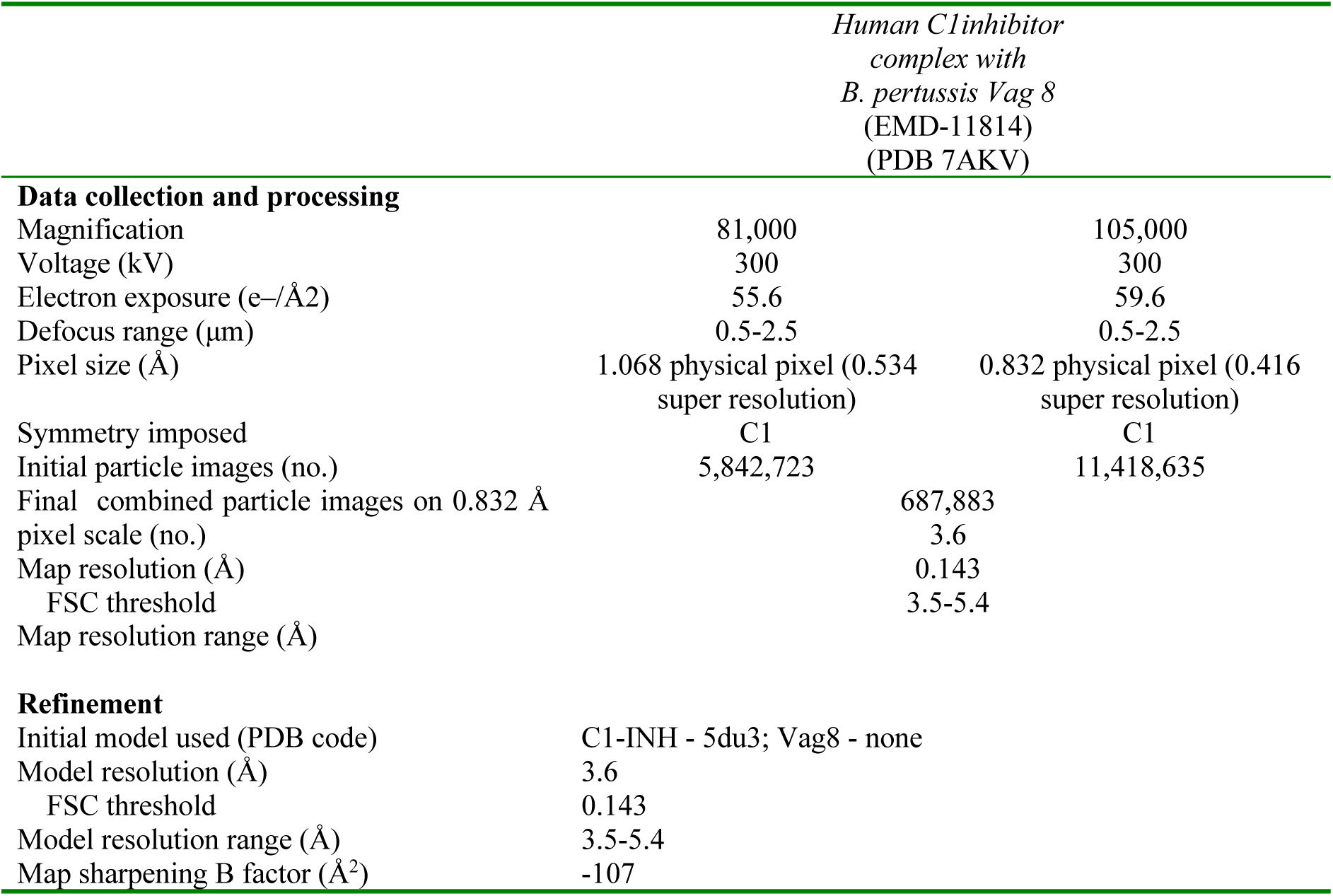
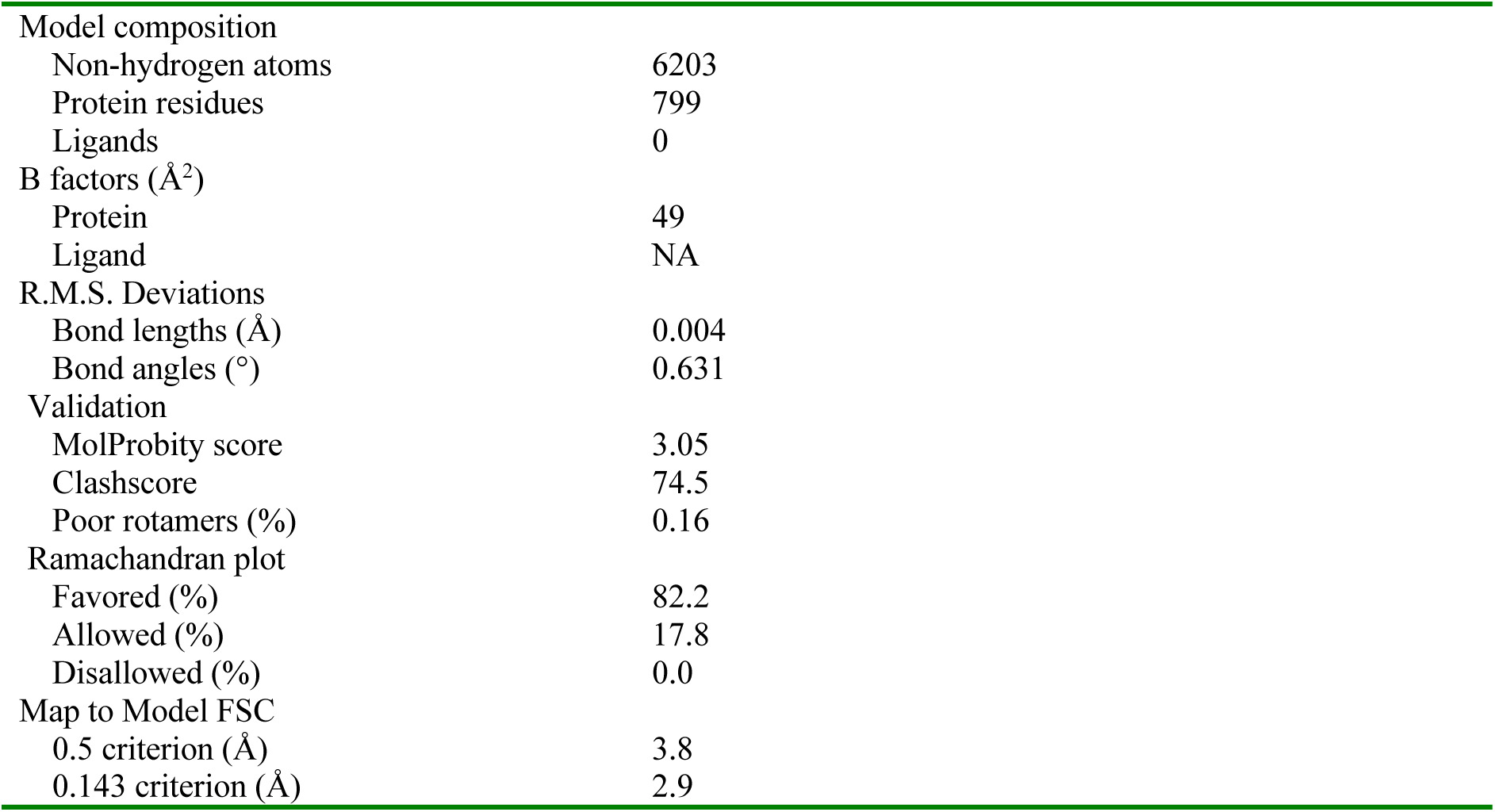
Structure solution and model quality

**Figure 2.**
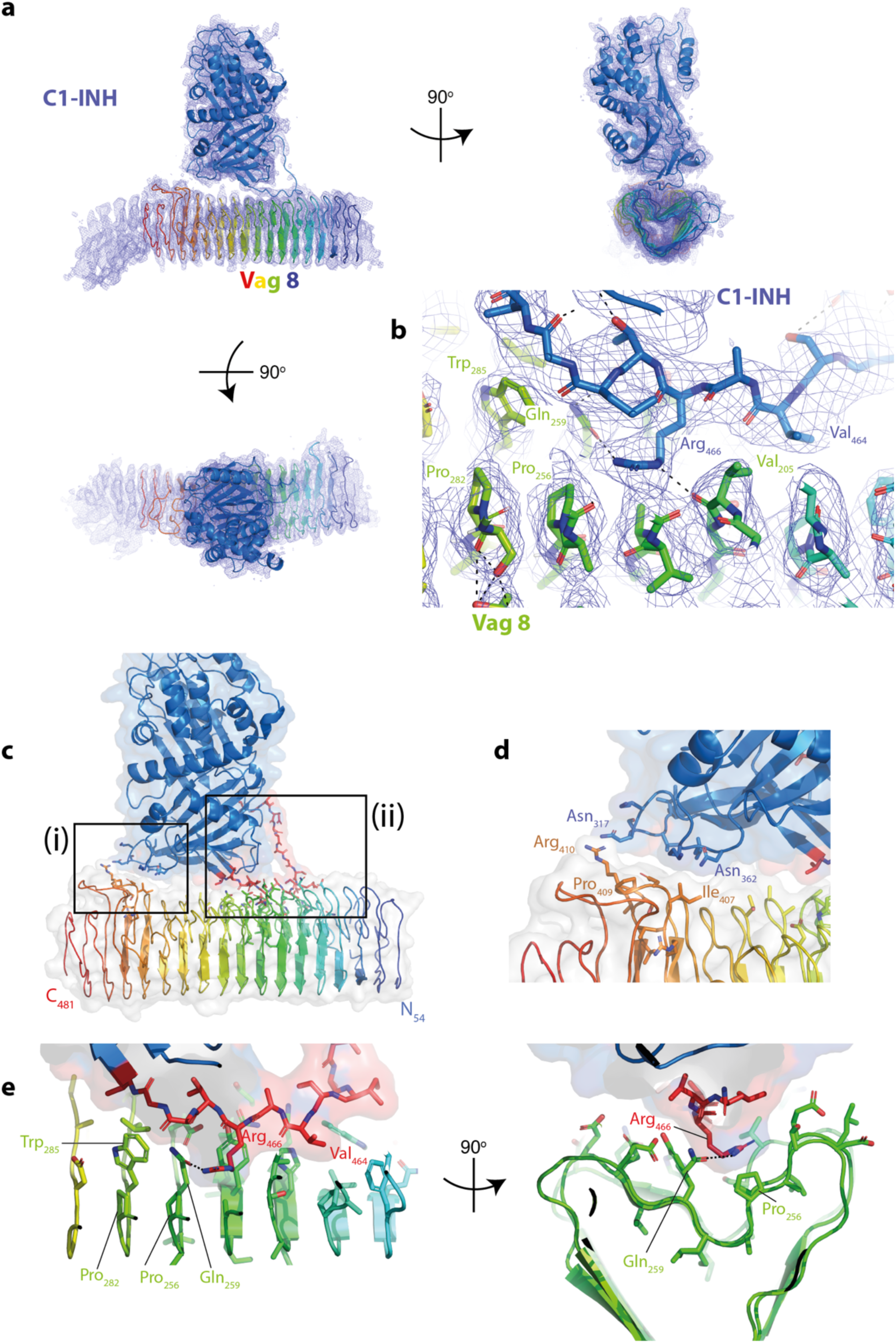
Structure of Vag8:C1-INH complex. (**a**)Views of the model of Vag8:C-INH in the experimental volume. Both proteins are shown in a cartoon representation, Vag8 coloured from blue at the N-terminus to red at the C-terminus and C1-INH coloured blue. Volume is contoured at 3s. Figure drawn using PyMOL (The PyMOL Molecular Graphics System, Version 2.0 Schrodinger, LLC) (**b**) shows a closeup of the central portion of the C-INH RCL in the Vag8 cleft with key residue interactions highlighted. (**c**) an overview of the complex with the two points of contact boxed (**d**) shows a closeup of the interactions in the smaller contact site boxed and labelled (i) in panel (**c**), (**e**) shows two views from the top and end of the complex of the larger interaction site boxed and labelled (ii) in panel (**c**).

The model for the complex reveals that C1-INH associates with the cleft within the Vag8 passenger domain beta-barrel, with two contact sites (Figure 2c). The first involves contacts between two loops at the base of the C1-INH Serpin domain (around residues 317 and 362) and one of the longer loops incorporated in the Vag8 beta barrel (residues 407-410) (Figure 2d). This is a fairly small contact area burying approximately 100 Å^2^ on each protein. The other point of contact is a much more significant interaction which buries the side chains of the majority of the RCL residues between 461 and 474 within the Vag8 beta-barrel cleft burying ∼600 Å^2^ on both components (Figure 2e).

To further probe the interactions seen in the complex we designed single and multiple point mutations in Vag8 to test their effect on complex formation. Mutant forms of Vag8 were expressed and purified then mixed with the C1-INH Serpin domain and complex formation was assayed by size-exclusion chromatography (Figure 3a, Table 2). With the exception of a mutation designed to sterically block binding of the RCL loop in the cleft by replacement of a small alanine side chain with a very large arginine side chain (A231R; Figure 3, Supplementary Table 1), mutation of multiple residues within the cleft to alanine was required to prevent formation of the complex emphasising the extended nature of the interaction site (Figures 2, 3).

**Table 2.** Binding analysis of Vag8 mutants by size exclusion chromatography. ‘+’ indicates Vag8:C1-INH complex peak seen, ‘-’ indicates proteins elute separately and no complex peak observed.

| Vag8 Mutation | Phenotype/C1-INH binding activity |
| --- | --- |
| H209A | + |
| A231R | - |
| Y234A | + |
| E237A | + |
| W285A | + |
| H209A Y234A E237A W285A | - |

**Figure 3.**
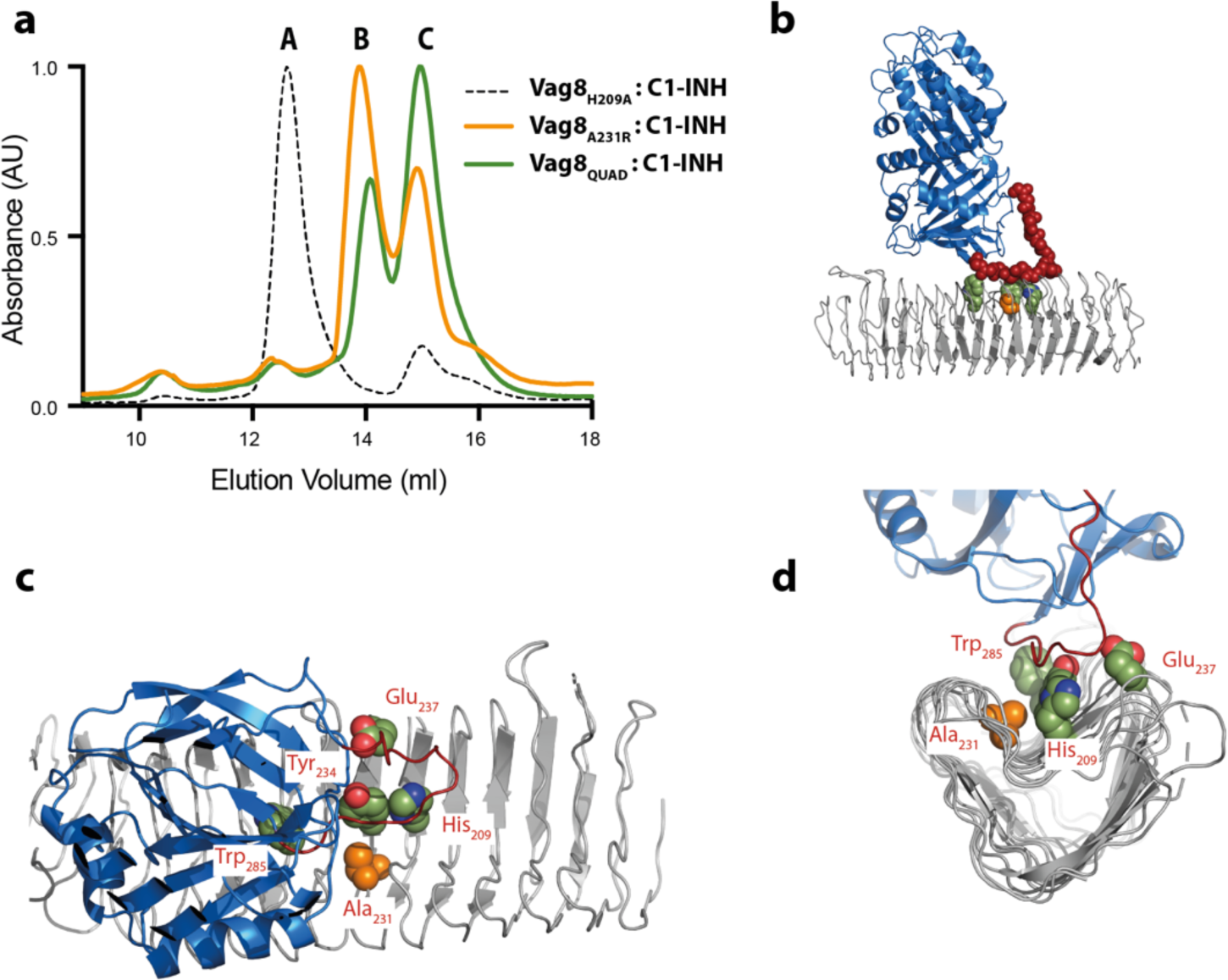
Mutation of residues within the interface abolishes complex formation. (**a**) 100µl of an approximately 1:2 molar ratio of Vag8 : C1-INH were mixed and purified using a S200 increase chromatography column. A,B and C denote the locations at which Vag8:C1-INH complex, Vag8 and C1-INH respectively elute. Vag8H209A (dashed black line) is one of the mutations that make up the Vag8QUAD set and is still capable of forming a complex with C1-INH (as are the other mutations that form the Vag8QUAD set in isolation, data not shown), whereas both Vag8QUAD (green) and Vag8A231R (orange) do not form any complex with C1-INH under these conditions and the two mixed components elute independently in peaks B and C (**b-d**) Views of the Vag8:C1-INH complex with Vag8 shown as grey cartoon, C1-INH as blue cartoon, residues mutated shown as space-filling spheres with carbons coloured to reflect colour scheme of panel (**a**). The C1-INH RCL loop is coloured dark red and the main chain atoms shown as spheres in panel (**b**). Panels (**b-d**) drawn using PyMOL (The PyMOL Molecular Graphics System, Version 2.0 Schrodinger, LLC)

## Discussion

*B. pertussis* targets regulation of immune, inflammatory and clotting processes by scavenging C1-INH using the passenger domain of Vag8. Our structure reveals that formation of this complex directly impacts on the physiological systems by masking the RCL required for C1-INH to fulfil its inhibitory activities within the bacterial protein. Unlike the native function of C1-INH, which results in formation of a covalent link between the RCL and the target, the inhibitor complex buries the RCL loop within the cleft of the bacterial protein via non-covalent interactions. Formation of a stable complex involving the RCL loop sterically occludes C1-INH interactions with its physiological partners.

*B. pertussis* is not the only organism that acts on these systems via scavenging C1-INH and it remains to be seen if other organisms use similar structural strategies to achieve inhibition.

## Materials and Methods

### Expression and purification of Vag8

Cloning of Vag8 passenger domain (residues 40-610) into a modified pRSETb plasmid has been reported previously (22). The recombinant plasmid was transformed into *Escherichia coli* C41 (DE3) cells which were then plated on LB-agar plates supplemented with 50µg/ml ampicillin. Protein production was carried by growing *E. coli* C41 (DE3) cells expressing Vag8pd in LB medium supplemented with 50µg/ml ampicillin at 37°C and 180 rpm until A600 reached 0.5-0.6. At this point, the culture was induced with 1 mM Isopropyl β-D-1-thiogalactopyranoside (IPTG) and further grown for 20 h at 24°C and 180 rpm. Cells were harvested by centrifugation at 5000 *g* for 10 min at 4°C. The cell pellet was resuspended in buffer A (50 mM Tris-HCl pH 8.0, 20 mM imidazole and 500 mM NaCl containing DNAase I and lysozyme). The cells were lysed using a Emulsiflex C5 homogeniser (Avestin) and the lysate cleared by centrifugation at 18000 *g*, 4°C for 45 min. The filtered supernatant was loaded onto a Ni-affinity chromatography column pre-equilibrated with buffer B (50 mM Tris-HCl pH 8.0, 20 mM imidazole and 500 mM NaCl). The Vag8 was eluted with a linear gradient of imidazole on an FPLC system (ÄKTA pure, GE Healthcare) using buffer B and buffer C (50 mM Tris-HCl pH 8.0, 500 mM imidazole and 500 mM NaCl). The eluted protein was dialysed overnight into buffer D (50 mM Tris-HCl pH 8.0 and 30 mM NaCl). The dialysed protein was subject to anion exchange chromatography and eluted by a linear gradient of NaCl using buffer D and buffer E (50 mM Tris-HCl pH 8.0, 1 M NaCl). Purified Vag8 was concentrated and the buffer was exchanged to buffer F (50 mM Tris-HCl pH 8.0, 150 mM NaCl) by ultrafiltration (Amicon Ultra, Merck-Millipore).

### Site-directed mutagenesis of Vag8

Single mutations in Vag8 (H209A, Y234A, E237A, and W285A) were introduced using Q5 Site-directed mutagenesis (NEB). The Vag8 quadruple mutant (H209A Y234A E237A W285A) was produced by Gibson Assembly of overlapping fragments containing the desired mutations using NEBuilder HiFi Master Mix (NEB). Purification of Vag8 mutants was done as described above for wild type Vag8.

### Expression and purification of C1-INH

A synthesised nucleotide fragment (codon optimised for *Saccharomyces cerevisiae*) encoding C1-INH amino acid residues 98-500 with Kozak and BiP signal sequence at 5′ end (GeneArt, ThermoScientific) was cloned using Gibson assembly (New England Biolabs) into pExpreS2-1 (ExpreS^2^ion Biotechnologies) plasmid, for protein production in *Drosophila* S2 cells, such that the mature recombinant protein had a His6 -tag followed by a 3C protease cleavage site at the N terminus. The recombinant plasmid was transfected into S2 cells following manufacturer’s protocol (ExpreS^2^ion Biotechnologies). Briefly, the recombinant plasmid was transfected into S2 cells and a stable cell line was selected over a period of four weeks while culturing the cells in EX-CELL420 medium (Sigma-Aldrich) supplemented with 10% (v/v) Fetal Bovine Serum (FBS) and 4 mg/ml zeocin. The stable cell line was maintained in EX-CELL420 medium supplemented with 10% (v/v) FBS, penicillin-streptomycin and amphotericin B, and cultured at 25°C, 110 rpm. For protein purification the stable cell line was split to a final cell density of 8 x 10^6^ cells /ml and cultured in EX-CELL420 medium, supplemented with penicillin-streptomycin and amphotericin B only, at 25°C, 110 rpm. The culture was centrifuged at 4500 *g*, 4°C for 30 min to collect the supernatant containing the recombinant protein four days after the split. The supernatant was filtered and incubated with His-tag purification resin (Roche) overnight at 4°C while mixing gently. The supernatant was then passed through a low pressure gravity flow column to collect the resin, which was then washed with buffer F. The protein was eluted using buffer G (50 mM Tris-HCl, pH 8.0, 150 mM NaCl, and 500 mM imidazole) followed by dialysis into buffer D. The dialysed protein was further purified using a MonoQ 10/30GL anion exchange chromatography column (GE Healthcare) by a linear gradient of NaCl with buffer D and buffer E. Purified C-INH protein was concentrated and the buffer exchanged to buffer F, 50 mM Tris-HCl pH 8.0, 150 mM NaCl by ultrafiltration (Amicon Ultra, Merck-Millipore).

### Preparation of C1-INH and Vag8 complex

The Vag8:C1-INH complex was prepared *in vitro* by incubating C1-INH in ∼1.5 molar excess with Vag8 at room temperature for 10 min followed by purification using size exclusion chromatography on a S200pg 16/600 column (GE Healthcare). The eluted fractions were analysed by SDS-PAGE followed by ultrafiltration to concentrate the protein complex.

### Size exclusion chromatography to assay the binding of Vag8 mutants to C1-INH

A 100 µL mixture of C1-INH (20 mM) and Vag8 WT or mutant (10 mM) was prepared at room temperature and injected onto a S200 Increase 10/300GL column pre-equilibrated with 50 Mm Tris-HCl 150 mM NaCl pH 8.0. The samples were eluted at 0.4 mL/min and 0.5 mL fractions were collected.

### Preparation of Cryo-EM grids

Four microliters of purified Vag8:C1-INH complex (0.5 mg/ml) was adsorbed to glow-discharged holey carbon-coated grids (Quantifoil 300 mesh, Au R1.2/1.3) for 10 s. Grids were then blotted for 3 s at 100% humidity at 8°C and frozen in liquid ethane using a Vitrobot Mark IV (FEI).

### Cryo-EM data collection, processing and analysis

Data were collected in counted super-resolution mode on a Titan Krios G3 (FEI) operating at 300 kV with a BioQuantum imaging filter (Gatan) and K3 direct detection camera (Gatan) using either a) a physical pixel size of 1.068 Å, a dose rate of 15 e−/pix/s, and an exposure of 4.23 s, corresponding to a total dose of 55.6 e−/Å^2^ or b) physical pixel size of 0.832 Å, a dose rate of 13.9 e−/pix/s, and an exposure of 2.97 s, corresponding to a total dose of 59.6 e−/Å^2^. All movies were collected over 40 fractions.

Motion correction, dose weighting, CTF estimation, particle picking and extraction were performed in streaming mode during collection using SIMPLE3.0 (42) as was 2D classification (42). *Ab intio* models were created in SIMPLE3.0 using particles selected from the chunks 1 & 2 further processing was performed in RELION 3.1 (43). The full workflow is described in Figure 1 but briefly, each data set underwent an initial round of 3D classification before 3D autorefine steps, beamtilt refinement, Bayesian polishing and further rounds of 3D classification (43). Chunks of data were combined as described in Figure 1 with the final volume calculated from 6 87,883 particles in C1. The resolution of the final volume is estimated as 3.6 based on FSC=0.143 criterion with the Local Resolution volume (calculated in RELION3.1 (43)) demonstrating that much of the core of the complex is at a resolution of 3.5 or better.

## Data availability

Coordinates and Volumes have been deposited in the PDB and EMDB respectively with accession codes 7KAV and 11814

## Acknowledgements

We thank the staff of the Central Oxford Structural Microscopy and Imaging Centre, Adam Costin and Errin Johnson and other members of the Lea group for assistance with various stages of the project. This work was funded by grants from the Wellcome Trust #219477, #209194 and #100298 and the Medical Research Council #S007474

## Author Contributions

AD cloned, expressed and purified complexes, performed binding studies. JCD prepared grids and collected single particle cryo EM data. EF screened cryo EM grids. DR cloned initial constructs. IJ, SJ & SML conceived study. AD, SJ & SML analysed data and wrote first draft of manuscript. SML processed cryoEM data and built models. All authors commented on final drafts of manuscript.

